# Neural trajectories improve motor precision

**DOI:** 10.1101/2025.07.01.662682

**Authors:** Wei-Hsien Lee, Xavier Scherschligt, Matthew Nishimoto, Adam G. Rouse

**Affiliations:** Neurosurgery Department, University of Kansas Medical Center, Kansas City, Kansas, USA; Department of Rehabilitation Medicine, University of Kansas Medical Center, Kansas City, Kansas, USA; Cell Biology and Physiology Department, University of Kansas Medical Center, Kansas City, Kansas, USA; Bioengineering Program, University of Kansas, Lawrence, Kansas City, USA; Electrical Engineering and Computer Science Department, University of Kansas, Lawrence, Kansas City, USA

## Abstract

Populations of neurons in motor cortex signal voluntary movement. Most classic neural encoding models and current brain-computer interface decoders assume individual neurons sum together along a neural dimension to represent movement features such as velocity or force. However, large population neural analyses continue to identify trajectories of neural activity evolving with time that traverse multiple dimensions. Explanations for these neural trajectories typically focus on how cortical circuits learn, organize, and implement movements. However, descriptions of how these neural trajectories might improve performance, and specifically motor precision, are lacking. In this study, we proposed and tested a computational model that highlights the role of neural trajectories, through the selective co-activation and selective timing of firing rates across the neural populations, for improving motor precision. Our model uses experimental results from a center-out reaching task as inspiration to create several physiologically realistic models for the neural encoding of movement. Using a recurrent neural network to simulate how a downstream population of neurons might receive such information, like the spinal cord and motor units, we show that movements are more accurate when neural information specific to the phase and/or amplitude of movement are incorporated across time instead of an instantaneous, linear tuning model. Our finding suggests that precise motor control arises from spatiotemporal recruitment of neural populations that create distinct neural trajectories. We anticipate our results will significantly impact not only how neural encoding of movement in motor cortex is described but also future understating for how brain networks communicate information for planning and executing movements. Our model also provides potential inspiration for how to incorporate selective activation across a neural population to improve future brain-computer interfaces.

**Summary:** Scientists have long studied how neurons in the brain help control movement. Traditionally, individual neurons were assumed to simply add together to create a signal to represent things like speed or force. But newer research shows that groups of neurons follow complex patterns over time—called neural trajectories. Here, we propose a computer model demonstrating how neural trajectories would enhance the precision of movement. Inspired by data from actual neurons recorded during a reaching task, we simulated neurons in motor areas of the brain and a downstream neural network to interpret and generate movement like might occur in our spinal cord and muscles. Movements were more accurate when neurons had different activations and timings relative to the desired movement rather than all working synchronously to generate a single signal. We conclude that the brain uses timing and coordination across many neurons— not just simple signals—to control movement. This work refines our understanding of how the brain signals movement and could improve technologies like brain-computer interfaces, which help people move or communicate using their thoughts.

## Introduction

The primary motor cortex (M1) contains neurons that modulate their firing rate to help orchestrate motor output through descending recruitment of muscles. Computational models that fit M1 neural activity to various behavioral outputs such as limb kinematics, muscle force, speed, and target location show reasonable performance in a subset of movements ^1,2^. Almost always, a linear summation of neural activity along certain neural dimensions is described as mapping to movement features. We believe this simplified readout assumption might be a key limitation to understanding how the brain represents movement, especially differences between large, fast movements and small, precise movements.

Point-to-point arm movements are executed with bell-shaped velocity profiles, characterized by smooth accelerations and decelerations ^3–5^. In addition to representing reach direction ^6^, Moran and colleagues ^7,8^ showed that individual neurons in M1 modulate their firing rates with movement speed, producing neural activations that closely mirrored the bell-shaped profile characteristic of reaching movements. This relationship has shaped the foundation of many neural decoding models, including those used in brain-computer interfaces (BCIs), where firing rates are typically modeled as encoding movement velocity — composed of direction and speed —as the primary variable to reconstruct motor intent ^9–11^. By extension, these linear models assume all the same neurons increase their firing proportionally when greater speed is required to reach further targets.

However, these linear models typically do not generalize well to diverse sets of movements, and actual neural activity has much richer spatial-temporal dynamics than could be explained by a simple readout of movement features from this neural activity ^12,13^. Several explanations have previously been given for why individual M1 neurons have diverse activations over time compared to the bell-shaped kinematic features. Spatiotemporal patterns of neural firing rates have been proposed to be due to the complex muscle activations needed for movement, involving agonists, antagonists, and fixators ^1,14^. Additionally, the motor cortex shows significant activity unrelated to movement, preparing for action in the ‘null space’ before muscle activity begins ^15,16^. Coordinated cortical neural signals across many neurons is also potentially crucial for motor control, learning, and adaptation, making motor signals robust to neuron loss ^2,17–20^. Despite the growing understanding of variance in neural encoding, all these models typically simplify downstream communication of these internal processes to a single neural dimension, using linear weighting and summation of firing rates for a degree-of-freedom of motor output ^14^.

The assumption that neural encoding can be reduced to a proportional output signal overlooks important aspects of neural recruitment. As the intended reach distance increases, the peak speed of movement increases ^21^ and greater force is required, which in turn demands the recruitment of additional muscle fibers. In the spinal cord, Henneman’s size principle ^22^ describes the orderly and serial activation of motor neurons based on cell size rather than proportional activation of all motor neurons. Smaller neurons drive slow-twitch fibers for lower-force movements, while larger neurons only engage fast-twitch fibers for higher-force demands. Similar patterns of selective recruitment, albeit less orderly, have been observed centrally: for instance, some pyramidal tract neurons in M1 fire more robustly during slow, precise movements than during fast, ballistic ones ^23^. More recently, cortical to motor unit recruitment has been shown to be functionally tuned depending on task conditions^24–26^. In addition to different recruitment patterns, motor cortex neurons also range in their timing with peaks in firing rate from 200-50 ms before peak hand velocity occurs ^7,27^. These combined findings suggest that increases in force or speed may not produce a uniform and proportional scaling of activity across the same cortical population, but rather selective co-activation and/or selective timing with varying subpopulations engaged. Rather than searching for a neural dimension that maps to an output degree-of-freedom, the range of neural geometries created by tuning and population-level dynamics should be considered ^28^.

Building on this perspective, we hypothesize the motor cortex does not simply scale firing rates within a fixed neural ensemble but utilizes selective recruitment to precisely execute different amplitude movements. We propose that selectively engaging subpopulations both in their co-activation and timing improves the estimation of reach distance in the presence of neural noise. To test this hypothesis, we analyzed behavioral and neural data from a non-human primate performing reaches to two different distances in the same direction. These initial findings reveal that reach distance modulates not just the magnitude of firing rates but varies the composition and timing of the active neural ensemble. Motivated by this observation, we developed a computational framework to investigate why the motor cortex may adopt this strategy. We provide evidence that selective co-activation of distinct neural subpopulations with different timings based on reach distance increases the separability between movement types in the neural population code. We demonstrate that varying both *which* neuronal subpopulations are engaged and *when* they are engaged provides the most accurate motor representation. This expanded representational space may allow brain-computer interfaces to decode movement amplitude and movement dynamics with a more precise prediction of motor intent.

## Results

### Distinct neural dimensions that encode reaching behavior

We analyzed neural firing rates when a monkey reached to two different distances recorded from M1 neurons across 13 sessions. We examined the neural subspaces that vary the most with given reach features of: i) velocity, ii) condition-invariant activity, and iii) reach distance, on reaches to two targets (near and far). Linear regression identified the two neural dimensions that best correlated with reach velocity (A-C). Demixed Principal Component Analysis (dPCA) identified neural dimensions with the most variance that were condition-invariant (D-F) and that best separated the two reach distances across time (G-I). By examining the three neural trajectories in their separate subspaces (C, F, I), distinct neural features are observed. The velocity representation shows neural firing rates correlated to the velocity of the reach as expected. However, we also observe two other distinct neural features that are less directly correlated with movement. The condition-invariant activity (F) shows that subsets of neurons are selectively active at different times, progressing through the two different neural dimensions. This activity is invariant between the near and far reach. The amplitude x time dimensions show neural dimensions that separate the two reach distances across time (I). Despite these being reaches in the same direction, the neurons are selectively co-activated in unique patterns for the two different distances from the very beginning of the movement. Notably, the amount of variance in the condition-invariant dimensions was larger than the velocity subspace identified by regression. The neural variance in the amplitude x time subspace was smaller than the velocity subspace but the near and far trajectories are most distinct in this subspace projection. Thus, these firing rate changes appear to not only be present but potentially provide additional information about the reaches to those dedicated to representing velocity. Together, these findings suggest that reaches are encoded across multiple neural dimensions—some dimensions separating the near versus far amplitude and others capturing temporal evolution—in addition to more classically described velocity tuned activity. These unique dimensions differ from a classical encoding model where a neural dimension representing force or velocity would carry direction and speed information together. Instead of most neural activity occurring in the top row of Fig. 1, additional condition-invariant and amplitude specific neural activity is present.

**Figure 1.**
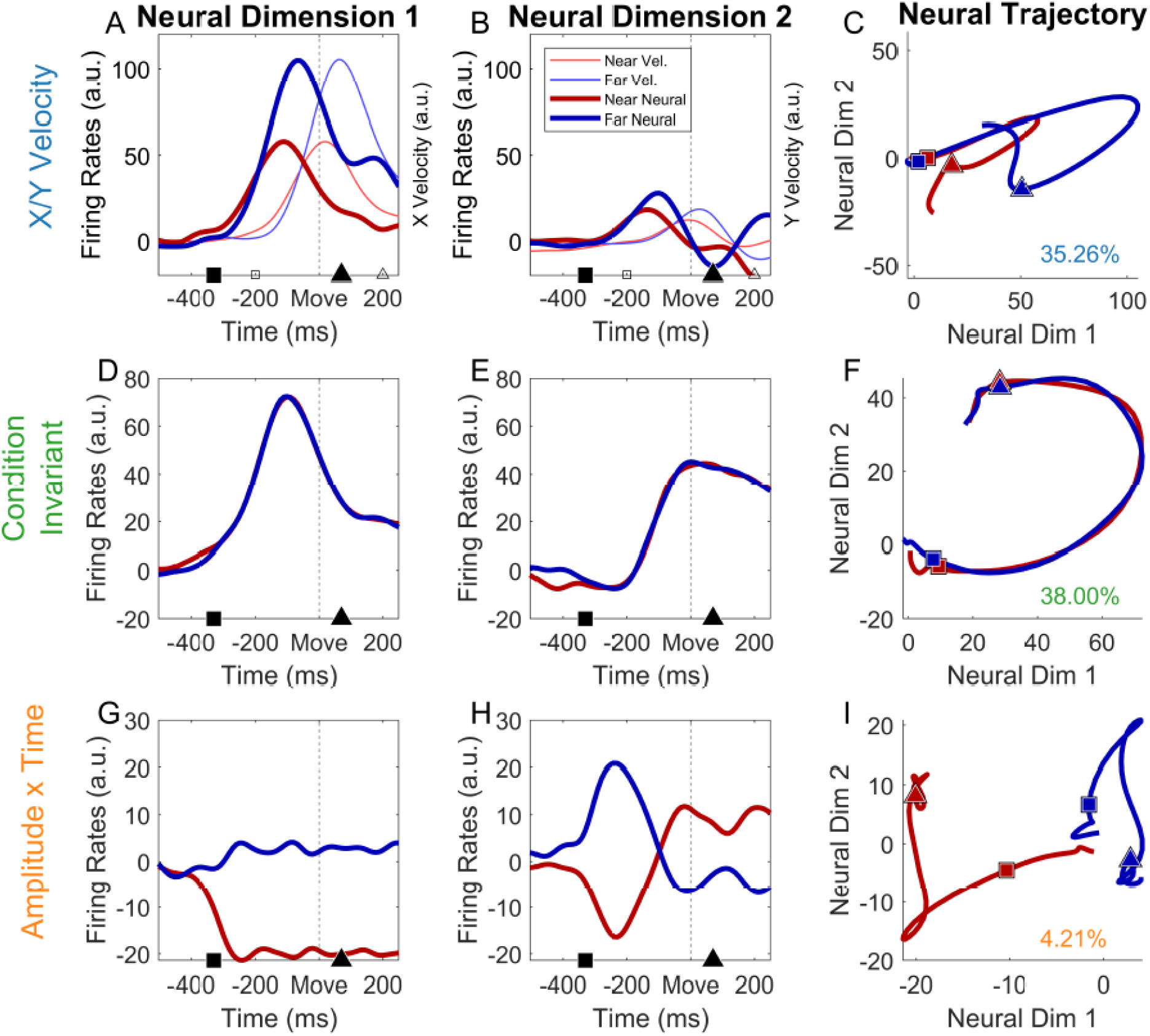
Neural subspace for reaches to two target distances. We applied a linear regression model for velocity and demixed Principal Component Analysis (dPCA) to neural population activity recorded during reaches to targets of two different distances to isolate and visualize behaviorally relative neural dynamics in three subspaces. <u>The X/Y velocity regression model identifies the neural spaces where velocity information is best predicted.</u> A) Neural dimension showing trial-averaged firing rates (bold lines) over time that best predicted X-velocity. The X-velocity is also shown (thin lines). Red lines indicate near targets and blue lines indicate far targets. The open square and triangle indicate the 200ms before and after the cursor left the center target, respectively. The highest correlation for neural activity with velocity occurred with a time offset of 130 ms, indicated with the filled symbols. B) Neural dimension showing trial-averaged firing rates over time best predicting Y-velocity. C) Neural trajectories best predicting velocity in the 2D neural space, first and second dimension indicate velocity prediction in X, Y reach dimensions respectively. <u>The Condition-Invariant activity identified with dPCA.</u> D) First neural dimension in the dPCA CI space, showing condition invariant average firing rate over time. E) Second neural dimension in the dPCA CI space, showing condition invariant average firing rate over time. F) Neural trajectories in the condition-invariant subspace combining D and E. <u>The Amplitude x Time dimensions identified with dPCA.</u> G) First neural dimension in the dPCA amplitude component space, showing amplitude dependent average firing rate over time for near and far reaches. H) First neural dimension in the dPCA amplitude × time interaction space, showing average firing rate over time for near and far reaches. I. Neural trajectories in the 2D dPCA space composed of G and H. Each subplots shows neural states from 500 ms before movement onset to 250 ms after leaving the target center. The time points of −330ms (square) to 70ms (circle) before leaving the center are shown on all plots to indicate the 130ms time lead of neural activity that best correlates with the velocity of movement during the −200:200ms window. All color-coded by red for the near target and blue for far target.

### Improved Decoding Performance with Higher-Dimensional Neuron Encoding Models

Using the condition-invariant and selective co-activation of neural activity as inspiration, we designed a neural simulation to test whether this type of neural activity can improve the accuracy of reaches. Synthetic firing rates were generated to encode reaches all in the same direction but for different distances. Four different neural encoding models of speed were used to generate the synthetic firing rates: (1) a proportional activation model; (2) a selective timing model with neurons activated differentially in the timing of their peak firing relative to peak movement speed; (3) a selective co-activation model created with preferentially activated neurons within distinct neural subpopulations; and (4) a combined model with both selective timing and co-activation. Fig. 2 shows the firing rates and neural trajectories in the neural subspace for the proposed neural encoding models.

**Figure 2.**
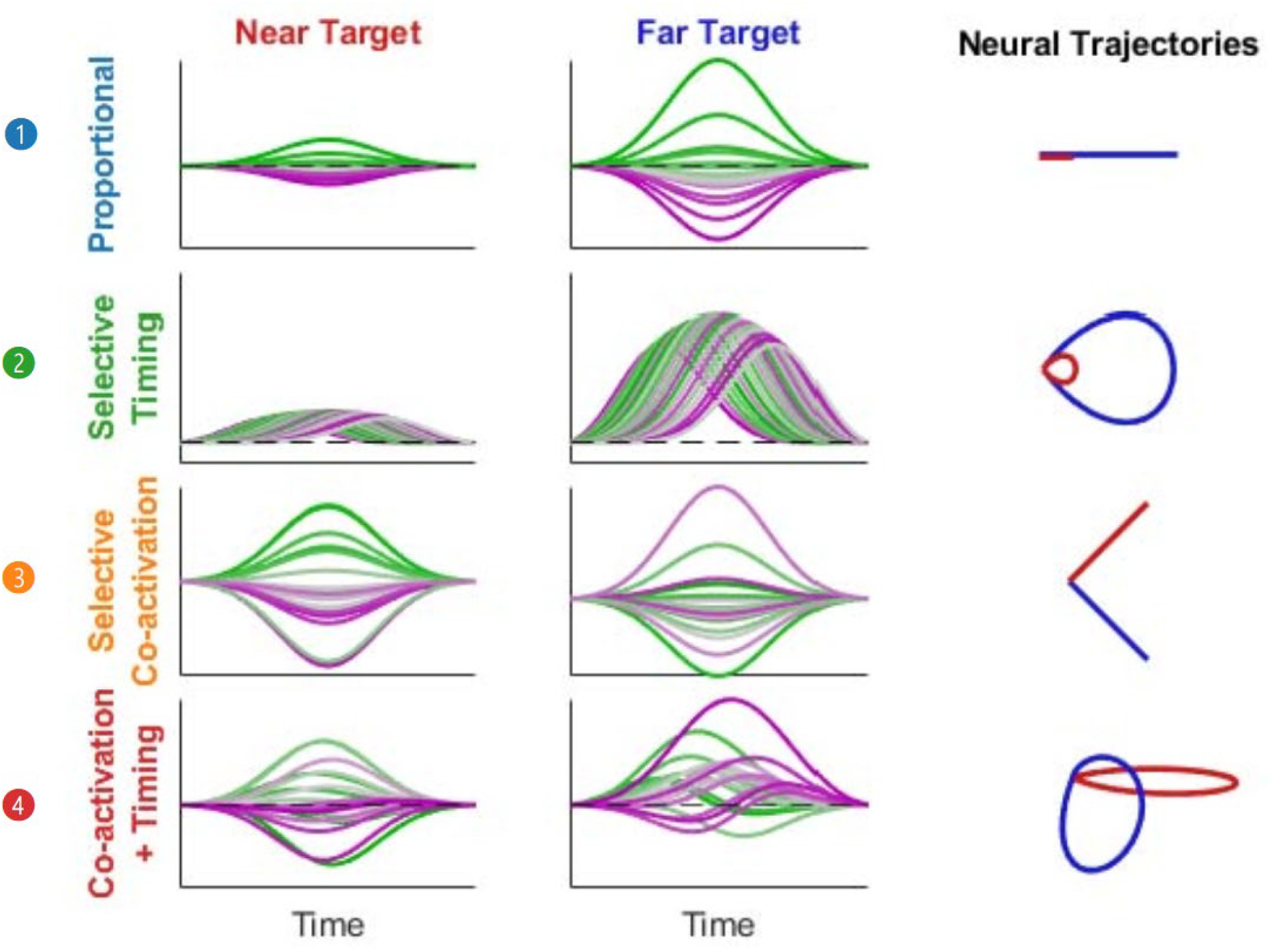
Firing rates and neural subspace for simulated encoding models. Firing rates of simulated neurons for 4 encoding models. The firing rates are shown for near (left) and far (right) distance reaches with velocity scaled to maintain the same reach duration. Model 1: In proportional activation, all neurons scale their firing rate in a linear relationship with velocity as assumed by a linear neural encoding model. Model 2: The selective timing of peak firing rates relative to peak speed is varied. The peak firing rates are proportional to speed for all neurons in this selective timing model. Model 3: Subpopulations of neurons are selectively co-activated with different peak firing rates depending on whether the target is near or far. All neurons have the same timing relative to movement. Model 4: A combined model includes both selective co-activation and timing for each neuron across the population. Neurons in each row are identified by the same color for near and far reaches.

Fig. 2, model 1, shows the proportional activation model where all neurons activate with a linear relationship to the distance of movement required, whether a near target (red) or far target (blue). This velocity tuning model utilizes a single neural dimension to represent movement along a single movement degree-of-freedom as often classically modeled in motor cortex. The magnitude along the single neural dimension indicates the speed along the single degree-of-freedom. Notably, this model creates neural trajectories that lay directly on top of each other for the different amplitude movements, as also observed in Fig. 1C for the start of the reach.

For our other encoding models, we added neural dynamics that occur in multiple dimensions. Model 2 utilizes selective timing of neurons (Fig. 2, model 2). The condition-invariant, cyclic neural activity we observed in Figure 1F inspired this model. By varying the lead and lag of various neurons, the current movement state, such as acceleration, peak velocity, or deceleration phases along those trajectories can be identified. While inspired by dPCA analysis of condition-invariant activity, our modeling adds variation in firing rate amplitude with reach velocity to the condition-invariant timing of firing rates. This introduces selective timing of neural activity into neural activity that also encodes the velocity of the intended reach. Model 3 utilized a selective co-activation encoding scheme for amplitude. The separate neural dimensions for the Amplitude x Time activity identified in Fig. 1I inspired this model. In the 2D neural space, the small amplitude movement occurs in one neural dimension while the largest movement occurs in an orthogonal, second dimension. Finally, we investigated model 4 incorporating both selective co-activation and timing which we hypothesize most closely mimics the various neural features naturally observed. Here, the neural trajectories are a summation of the 2) co-activation and 3) timing models with their mixture encoding the movement information.

We trained a similarly structured MLP (Multilayer Perceptron) – LSTM (Long-short term Memory) Recurrent Neural Network (RNN) mixture model for each encoding model to provide movement decoding (described in methods). We used the same decoding architecture to directly compare performance across the four encoding models. The same white noise levels were added to each synthetic neural encoding models.

### Comparable Performance Improvements with Selective Co-activation and Selective Timing

#### Models

The results of our simulation, summarized in Fig. 3, show that both the selective co-activation and selective timing neural encoding models lead to improved decoding of movement velocity compared to the proportional model (blue line). In particular, the selective co-activation model (orange line), where distinct subpopulations of neurons are preferentially activated with varying peak firing rates depending on whether the target is near or far, significantly enhances decoding performance across all tested noise levels. A similar improvement was also observed when using a temporal progression within the neural dynamics to signal the phase of movement. Better performance also occurred for the selective timing model (green line) compared to the proportional activation model (blue line). In this scenario, the decoder presumably uses the current neural trajectory location to identify the phase of movement-acceleration or deceleration- and better integrates the decoded velocity across time to more accurately predict movement, highlighting how the precise timing of neural activity can lead to increased motor precision.

**Figure 3.**
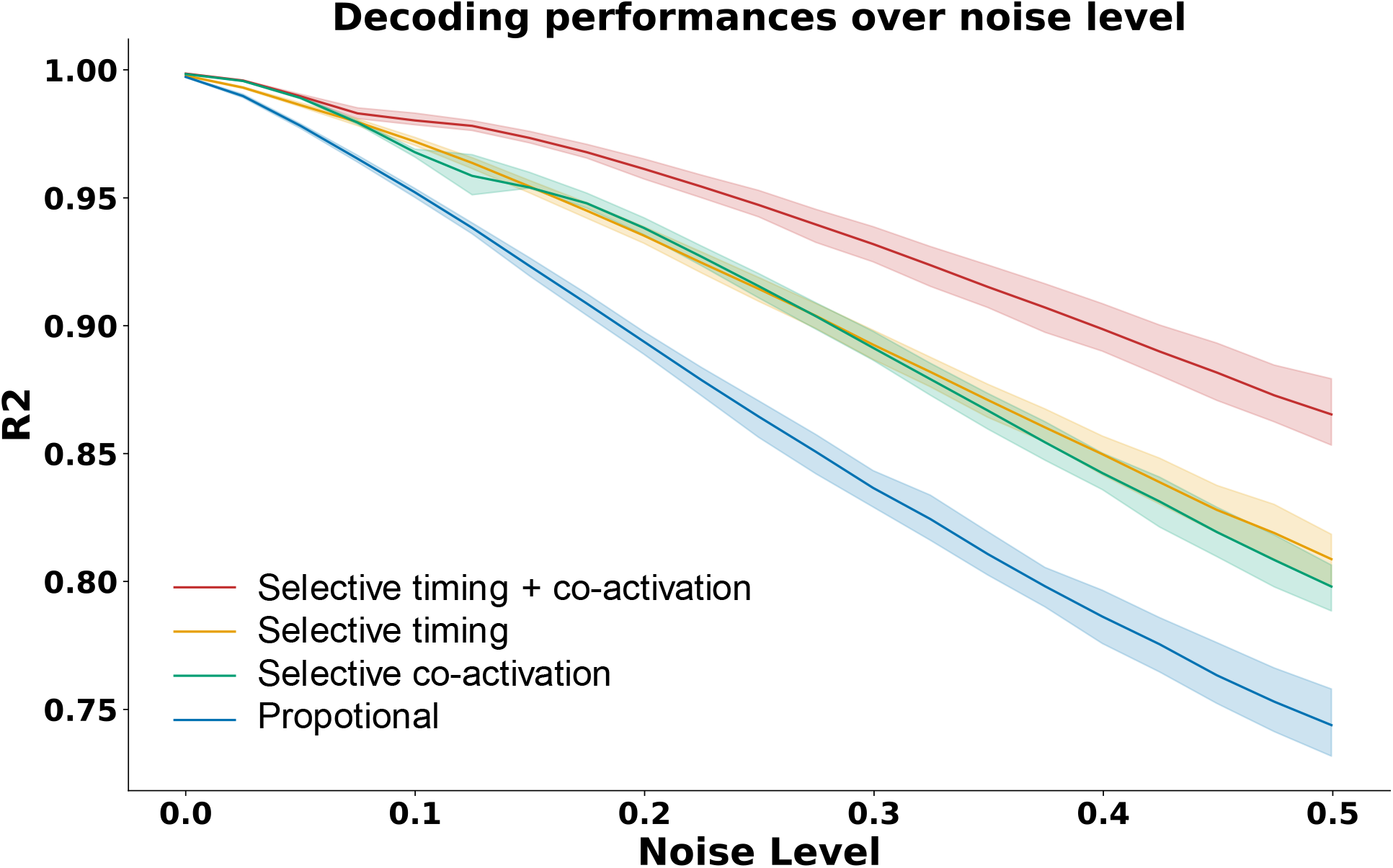
R2 performance plot. The correlation coefficient (*R*^2^) indicates the decoding performance of the propose encoding models, proportional model (red), selective co-activation model (orange), selective timing model (green), selective timing + co-activation model (red), over different noise levels. The average R^2^ of the model’s performance is color-coded in bold color lines and the 25-75 percentile range of performances over bootstrapping is in shaded area with corresponding color keys. The proportional model was significantly worse at all noise levels (Cliffs’ delta > 0.75). Selective timing + co-activation model was better than all other models (Cliffs’ delta > 0.75) except for being equivalent to selective co-activation at low noise levels (σ= 0-0.075). selective timing and selective co-activation model were also equivalent (Cliffs’ delta < 0.75) at low noise levels (σ= 0-0.05).

We next examined whether the selective timing model outperforms selective co-activation model or vice versa. Here we compared the decoding performance between selective timing (green line) and selective co-activation model (orange line) in Fig. 3. The results revealed no significant performance differences (Cliff’s delta < 0.75) between the selective timing model and the selective co-activation models when there was moderate noise or above (σ>0.075), suggesting similar performance improvements are possible with either encoding scheme.

### Encoding both spatial and temporal information provides the best neural representation of movement

Finally, we examined the encoding scheme when selective timing and co-activation is combined. This combined model has neural trajectories with a mixture of amplitude and timing to embed movement information. Similar to the selective co-activation encoding scheme, the angle of the trajectories indicates the speed of movement. Additionally, the cyclic pattern within each trajectory from the varying timing of firing rates allows the movement phase to be identified. Together, both selective timing and co-activation enable a spatiotemporal encoding. As shown in Fig. 3, the selective timing + co-activation model provides decoding performance better than the other three proposed models (Cliffs delta > 0.75) except at very low noise levels (σ < 0.075).

### Similar tradeoff between selective co-activation and timing

To better quantify the effect on encoding quality of selective timing, co-activation, or both, we analyzed a parameter sweep across varying differences in timings and co-activations as shown in Fig. 4. The parameter sweep was performed under a fixed noise level (*σ* = 0.3) within the encoding model. The decoding performance (R^2^) is presented as a heatmap across the parameters. The darker blue indicates better decoding performance under that given parameter. The result shows an improvement in decoding performance as both timing differences and neural co-activation differences increase. Here, we define the timing difference as the time difference between the earliest and latest firing neurons (in ms). The co-activation dimensionality difference is changed by altering the angle between two representational neural dimensions within a 2D neural manifold ^29^. The co-activation ranges between 0 to 90 degrees to indicate how much the two representational neural subgroups overlap with each other between a 1D (fully overlapped with neural variance in a single dimension) and 2D (orthogonal to each other, utilizing 2 neural dimension) representation, respectively. We observe the effects of selective timing and co-activation are synergistic. In general, only a small amount of difference in co-activation or timing is needed to realize much of the performance improvement when the other feature difference is large.

**Figure 4.**
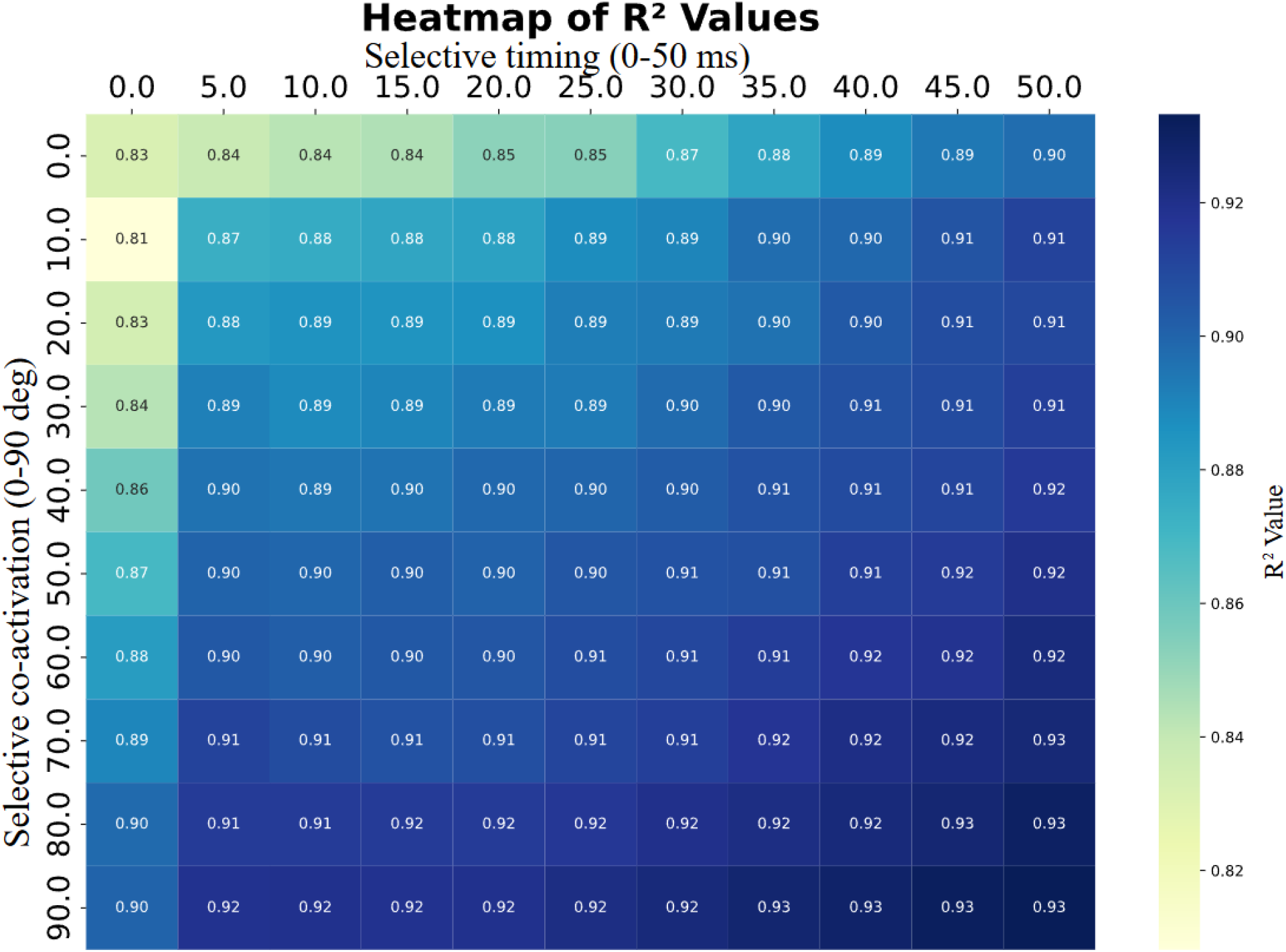
Selective timing/co-activation model parameter sweep analysis on a fixed noise level. A heatmap of the parameter sweep performed across timing and co-activation levels, quantifying the improvement in encoding both dimensional and phase information at various level. The proportion of neuron co-activation that’s encoded in the sweep is determined by degrees. 0 degree indicates no separate dimensional encoding of movement amplitude, and 90 degree indicates two orthogonal dimensions of the neural space to encode movement amplitude. The selective timing in the sweep is determined by time lag in milliseconds. 0 ms indicates no timing difference across neurons, while higher values up to 50 ms reflect the time lag between the earliest and latest firing neuron, capturing the temporal spread of activation across the population. Darker blue color indicates a better performance in the model. The parameter sweep is performed on encoding models at a single noise level (*σ* = 0.3).

## Discussion

Our results underscore the critical role of selective recruitment of neurons in precisely encoding movement. The proposed selective co-activation and timing models demonstrate significantly improved decoding performance compared to a proportional activation model for different reach speeds. By including the selective information of neuron co-activation, the resulting neural trajectories become more distinct, allowing for better separation of movement types. This idea has some similarities to the idea of motor cortex reducing “tangling” of neural trajectories and BCI decoders that rely on distinct states formed by neural factor trajectories, so different movements are unambiguously represented in the dynamic neural system ^30,31^. These ideas have previously been applied to separating distinct tasks and the challenge of generalization between them. Here, the advantage of these separated neural features is evident where movements of varying speeds need to be differentiated even within a single reach direction. Our decoder could predict the upcoming peak speed at an early stage by analyzing the initial neural trajectory leaving the origin (idle state) before the neural activity peaks in its depth of modulation (the furthest neural state from the origin) (Fig. 2). Subsequently, the higher-dimensional models provide bigger neural differences between near and far conditions, facilitating a more robust representation of neural activity resilient to noise.

The idea of neural trajectory encoding also shares similarities to the idea of movement fragments presented by Hatsopoulos and colleagues ^32–34^ Like us, their work identified the importance of time-varying tuning of individual neurons to upcoming movement. It also recognized the importance of holistic submovements rather than thinking of cortex representing upcoming movement at a single instance of time. Their model, however, focused more on how a single neuron might have a preferred, often curved, movement trajectory with different directions at various future timepoints. Our work focuses on how multiple neural dimensions might better represent upcoming movements—even straight-line reaches— more accurately.

Our results highlight the utility of multiple neural dimensions for improving decoding performance. Our selective co-activation model performs comparably to the selective timing model, suggesting that either temporal variation or specific neural co-activations can enhance decoding. The combined encoding model, which combines both spatial and temporal features, emerges as the most comprehensive representation of movement. By integrating selective timing and co-activation information into higher dimensional neural spaces, this model adds a critical layer of precision, allowing for the differentiation of movement states such as acceleration, peak velocity, and deceleration. Again, this makes it easier to predict both speed and movement states even at movement initiation. The neural trajectories enable a more detailed representation of the temporal progression of movement besides the instantaneous speed. These findings align with our hypothesis that natural brain encoding leverages both spatial and temporal neural features to improve movement representation.

Our neural trajectory results also provide a tantalizing result when considering not only motor output but interarea brain communication. Key concepts of the cerebellum are internal models and an efference copy leading to a teaching signal ^35–37^. While this teaching signal is often conceived separate from a motor output signal, a predictable neural trajectory has the beneficial property that the early phase of the trajectory signals upcoming movement. This combined representation of current and future movement could provide information to cerebellum circuits about how an executed movement compares to the original motor plan. Mixed selectivity in cortical neurons may also facilitate communication across brain regions by embedding contextual and motor variables into high-dimensional representations, enabling downstream structures to flexibly decode and compare motor plans ^38^. The initiation of several simultaneous neural trajectories could represent possible motor actions that can then be either reinforced for execution or suppressed to select a single desired motor action. Thus, motor action selection by the basal ganglia ^39,40^and other brain structures that gate motor output might be aided by neural trajectories.

While our findings highlight the advantages of high-dimensional encoding models, several limitations remain. Artificial neural networks can uncover hidden structure and predict movement states with high accuracy, but often lack biophysical realism. Our results suggest that the selective co-activation and temporal encoding schemes may be equally effective and potentially integrated within the neural system, pointing to a gap in understanding for when selectively activated neurons and temporal information might be utilized to optimize movement control. The combined encoding model suggests that these strategies are synergistic in our specific model. Although the combined model points to synergy, further studies are needed to quantify the relative contributions of these strategies. In addition, our simplified trajectory-based model, while advantageous for accuracy and communication, constrains online correction and may not generalize to naturalistic, noisy environments. Future work should refine neural network models for interpretability, enabling direct comparison with biological mechanisms, and explore how spatial and temporal encoding are integrated to support adaptive motor control. Moreover, further exploration of higher-dimensional selective co-activation and timing encoding schemes could reveal whether and how the brain leverages both strategies synergistically.

## Methods

### Behavior task

We utilized a Center-Out Amplitude (COT Amplitude) task to characterize neural activity correlated to reaches to two different distances. A male, six year old 10 kg rhesus macaque participated in this study. All animal care and research procedures adhered to the Guide for the Care and Use of Laboratory Animals. Approval was granted by the Institutional Animal Care and Use Committee at the University of Kansas Medical Center.

The monkey was trained to perform the COT Amplitude task using an exoskeleton robotic arm to control a cursor on a 47’’ LED display provided by NHP KinArm (BKIN Technologies, Kingston, ON). The animal was body-restrained in the KinArm chair, as well as forearm and elbow restrained on the exoskeleton arm. The exoskeleton arm could move freely at the elbow and shoulder on a 2-Dimensional plane parallel to the ground in this task. The cursor on the screen represented the center of hand position in the exoskeleton arm which can freely move within a 1000-by-1000 pixel workspace. A circular cursor indicated a single point in the workspace. Dexterit-E data acquisition and experimental control was customized within a Simulink software environment to record the cursor position and trial event information and was down sampled to 100 Hz for analysis with the neural data.

We recorded 13 sessions of neural recordings in a time span of 1 month. The monkey successfully completed a total of 8867 trials across the recording sessions. During the COT Amplitude task, the monkey was instructed to perform a center-out reach to 1 of the 8 peripheral targets in four reaching directions equally spaces by 90^°^ around the origin after obtaining and holding the cursor at the center target for 300-500 ms with a radius of 1.25 cm. Each of the reach directions includes two targets with different reach distances, a closer target and a further target. In this study, we are only using neural and behavioral data from one of the four directions to show the different encoding schemes between the two reaching distances to simplify and align with the proposed hypothesis and simulation models. All neural data processing were done with a custom, automated spike-sorting pipeline and used in the following population analysis ^41^.

### Linear regression

We implemented linear regression to examine how neural firing rates evolved in relation to cursor movement X and Y velocity during one direction of reaching in COT Amplitude task. To begin, we extracted trial-averaged firing rates from 1,228 neurons during near and far reaches and aligned them with the corresponding conditions. We time aligned the data to the when the cursor left the center target for each trial. Principal component analysis (PCA) was performed on the trial-averaged firing rates within a time window spanning from 300 ms before to 300 ms after leaving the target center. This reduced the data to 15 neural dimensions, capturing dominant features of population activity across time and movement contexts.

To link neural dynamics with behavior, we next fit a linear regression model that predicted X and Y velocity based on the top 15 neural principal components. We predicted the kinematic data within the time window 200 ms before until 200 ms after leaving the center target. We used a sliding window from 200-0ms of neural activity preceding movement and identified that neural activity 130ms before movement best correlated with movement velocity. This allowed us to assess how well low-dimensional neural trajectories captured the velocity of the movement. By using this behavioral variable to identify the two best correlated dimensions in the neural state space, we visualized how motor parameters were encoded over time within the population activity. This approach revealed behaviorally tuned dimensions in the neural state space that reflected continuous encoding of movement features.

### dPCA analysis

We applied demixed principal component analysis (dPCA) to identify and emphasize the key features of complex population activity ^42^. This method decomposes the population activity into a limited number of demixed components that capture the most variance in the data related to the specified task features such as time and reach distance.

We again applied principal component analysis (PCA) to reduce the dimensionality of the firing rate data to 15 dimensions. The PCA scores were then reorganized into a structured format suitable for dPCA, preserving the original data dimensions while enabling task-specific decomposition. The neural data were arranged into a tensor of size N×T×A, where N represents the number of neural PCs, T time points (from 300 ms before until 300 ms after leaving the center targe), and A amplitudes. This transformation allowed dPCA to systematically separate neural dimensions that encode task parameters, distinguishing between condition-dependent neural dynamics that vary with time.

Once dPCA was applied, the projected neural trajectories were then computed by multiplying the PCA-transformed neural activity with the dPCA demixing weights. We analyzed neural population activity in these distinct component spaces derived from dPCA, focusing on three key representations: (1) condition-invariant time-varying neural dynamics, reflecting the shared temporal structure of neural activity across the two reach distances; (2) condition-dependent representations, where neural activity was modulated by reach distance, highlighting task-specific encoding; and (3) time × amplitude interactions, which revealed how reach distance influenced neural dynamics over time.

### Neural simulation

In the study, we simulated motor cortical activity to generate movements. The simulated task mimicked the kinematics of a single direction reaching task from movement onset to target acquisition. A bell-shaped speed profile like those observed in natural point-to-point reaches was used ^43^ without considering other factors such as position or acceleration.

We simulated 1000 reaching trials, each with a randomly assigned velocity peak within a range of 0.25 to 1 arbitrary unit. Each trial had a duration of 200 ms with sampling of 5 ms. Despite various magnitudes of the movements, the speed profiles across all trials shared the same symmetric gaussian shape with a standard deviation of 33.3 ms. Example movement speed profiles are shown in Figure 1 with the i) red curve representing a small magnitude reaching movement with lower speed and ii) blue curve for a large magnitude reaching movement with high speed. In our simplified task, all movements were the same duration with the speed amplitude directly related to the distance reached. These same 1000 simulated reaching trials were used for all the various simulated neural models.

### Speed encoding models

We designed four neural encoding models of speed to describe the neural activity based on selective activation of neurons: (1) a **proportional activation** model, in which all neurons scale their firing rates linearly with movement velocity; (2) a **selective timing** model, in which all neurons encode speed proportionally, but differ in the timing of their peak firing relative to peak movement speed; (3) a **selective co-activation** model, where distinct neural subpopulations are preferentially activated depending on movement amplitude (near vs. far), with all neurons peaking at the same time relative to movement; and (4) a **selective timing + co-activation** model that includes both selective co-activation and timing for each neuron across the subpopulation. The toy neuron models were tuned solely to velocity for simplicity. The same Gaussian white noise was added to the firing rates of all neural models. In the proportional activation model, the firing rate was generated by multiplying the actual speed vector and the speed tuning weights of each neuron with following equation:

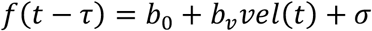

Where *f* is the neural firing rate; *b*_0_ is the baseline firing rate; *b*_*v*_ is the speed encoding weight vector across the neurons; *vel* is the actual movement speed across each trial.

The selective timing model introduces a change in time lags for the different neurons tuning to speed. We manipulated the shift of the time across the neurons to create a population code with the same velocity shape as other models to represent the same velocity profile. The neural activity would then be described as follow

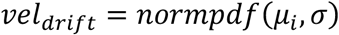

Where *μ*_*i*_ ranges from 0 to *t*_*max*_ ms to define the time lag for each neuron *i*. For the main simulation of model 3 (and model 4), *t*_*max*_ = 50 ms. For the parameter sweep in Fig. 4, *t*_*max*_ ranges from 0 to 50 ms.

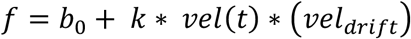

*vel(t)* applies a bell-shaped envelope to match the overall speed profile and ensure all neural firing rates begin and end at baseline. A scale factor, *k*, was used to normalize the total variance across all neural firing rates between Model 1 and Model 2 to provide a fair comparison in the simulation.

In the selective co-activation model, we used two orthogonal neural dimensions of tuning weights to encode speed. These two vectors combine together to generate the neural activity as follows:

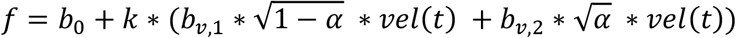

Now, *b*_*v*,1_ and *b*_*v*,2_ define the two weight vectors across the neurons creating two neural dimensions of encoding. In the main simulation of model 3 (and model 4), *b*_*v*,1_ and *b*_*v*,2_ are orthogonal. For the parameter sweep presented in Fig. 4, the angle between *b*_*v*,1_ and *b*_*v*,2_ was varied between 0 to 90 degrees to range from 1D encoding (fully aligned as in Model 1) to orthogonal 2D encoding (as in Model 3). The *α* value is a scaling term that is 0 for the smallest speed movement possible and 1 for the largest speed movements that shifts the encoding from the *b*_*v*,1_ dimension for small movements to the *b*_*v*,2_ for the largest movements. Again, a scale factor, *k*, was used to normalize the total variance across all neural firing rates between Model 1 and Model 3.

Finally, we tested a combination model with a linear sum of both selective co-activation and timing model with various weights defined for both the amplitude and phase encoding as described above.

### RNN decoder design

We employed an RNN mixture model which integrates a MLP and a LSTM network to train on the hypothesized four encoding models of neural activities and used to evaluate decoding performance from those models.

The MLP serves as an initial feature transformation stage, mapping the input neural activity (N=20 dimensions) through nonlinear combinations into the same dimensional space. This preprocessing step allows the network to learn more structured representations of the neural data before passing them to the subsequent LSTM for temporal modeling..It consists of two fully connected layers with ReLU activation functions, designed to capture the non-linear relationship in the raw neural data. The last layer readout in MLP, preserving the dimensions of neural data, is then passed down to the subsequent LSTM network as input. The non-linear regression process within the hidden layers of MLP can be described as follow.

Let *x* ∈ ℝ^*n*^ represent the input neural data. The MLP consists of *L* layers, each defined by a set of weights *W*^(*l*)^ and biases *b*^(*l*)^ for layer *l*, where *l* ∈ {1,2, …, *L*}.

The non-linear transformation at each layer can be described as:

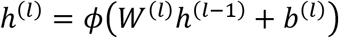

where:

- *h*^(0)^ = *x*,
- *h*^(*l*)^ is the output of the *l*-th layer,
- *ϕ*(⋅) is non-linear activation function ReLU (Rectified Linear Unit), defined as *ϕ*(*z*) = max(0, *z*).

The final output of the MLP, *h*^(*L*)^, provides the encoded representation of the neural data, which is then passed to the LSTM network.

The LSTM network comprises two hidden layers with tanh activation functions and sigmoid gates to manage the flow of information across time steps and finally infer the speed over time steps at the output. The LSTM then processes the MLP-encoded inputs (N=20) to infer the behavioral output, producing a one-dimensional estimate of movement speed at each time step.

We optimized the decoding model’s hyperparameters by conducting a comprehensive sweep over weights and biases. This process involved systematically varying parameters including learning rate, number of units per layer, regularization coefficients, and initialization methods. Performance metrics, including accuracy and loss, were used to evaluate each configuration. The optimal hyperparameter set was selected based on its ability to enhance the capacity to capture temporal dynamics in the neural data, resulting in improved predictive performance.

### Quantification of Decoding performance

We used the correlation of determination (*R*^2^) to evaluate how well velocity can be predicted using the same deep learning architecture across different encoding models. A single-trial *R*^2^ is calculated to assess the performance of the MLP-LSTM model on individual trials. This metric provides a detailed measure of the model’s predictive accuracy, showing how effectively it captures the relationship between neural activity and movement speed within each trial. By examining single-trial *R*^2^, we gain insight into the model’s ability to decode speed from neural activity and account for variability observed in single-trial movement patterns.

To evaluate overall performance, a trial-averaged *R*^2^ is calculated by averaging the *R*^2^ values across all trials for each prediction made by the MLP-LSTM model. This trial-averaged *R*^2^ provides a comprehensive measure of the model’s generalization capability, reducing the influence of noise and trial-to-trial variability. By aggregating the *R*^2^ values, we emphasize the predictive power of each encoding model under varying conditions, highlighting their ability to capture the relationship between neural activity and movement dynamics effectively.

### Statistics

We performed multiple iterations using random neural tunings in the encoding model and randomized initial conditions in the similarly structured MLP-LSTM decoding architecture to test for repeatability and robustness of the observed model differences. The white neural noise in the encoding model was also randomly generated but the same noise instance was used across the four encoding neural models to minimize the effect of randomized noise on model differences. Models were trained through the similarly structured MLP-LSTM decoding architecture to get a confidential interval of the performance difference between those encoding models and identify outliers during training.

We used Wilcoxon signed-rank test as a non-parametric statistical tool to conduct pairwise comparison between decoding performance of the four encoding models. As is common with computational models with many iterations, the Wilcoxon test showed statistically significant mean differences in decoding performance (p<<0.001) between all pairs of models at each noise level. We then chose to calculate the Cliff’s delta values for each comparison groups to test for meaningful effect size differences in decoding performance. We chose a Cliff’s delta greater than 0.75 as a meaningful difference in effect size. This corresponds to one model outperforming the other model 75% of the time. We also present the 25 to 75 percentiles for R^2^ values across trials, allowing us to quantify the variability and reliability of the model predictions.

### Notation

|  |  |
| --- | --- |
| $f$ | neural firing rate |
| $b_0$ | baseline firing rate |
| $b_v$ | speed encoding weight vector across the neurons |
| $vel$ | actual movement speed across each trial |
| $vel_{drift}$ | describe the time lag of firing for each neuron |
| $h^{(l)}$ | output of the $l$-th layer in MLP network |
| $\phi(\cdot)$ | non-linear activation function ReLU |

## Acknowledgements

We wish to thank Luke Andriano, Evan Schrader, and Lauren Wynn for their assistance with data collection.

## Data and code availability

The code used for the simulation is publicly available through *github* at: https://github.com/Precision-Neural-Dynamics-Lab-KUMC/Neural-Speed-Encoding-Model

The neural and behavioral data that support the findings of this study are available from the corresponding authors upon request.

## Notes

https://github.com/Precision-Neural-Dynamics-Lab-KUMC/Neural-Speed-Encoding-Model

### Competing Interest Statement

The authors have declared no competing interest.

### Summary of Updates

Minor edits to descriptions of methods and editing for improved clarity

### Funding

- NIH NINDS: R00NS101127

